# Profile of *Mycobacterium tuberculosis*-specific CD4 T cells at the site of disease and blood in pericardial tuberculosis

**DOI:** 10.1101/2022.05.12.491749

**Authors:** Elsa Du Bruyn, Sheena Ruzive, Patrick Howlett, Ashley J. Jacobs, Cecilia S. Lindestam Arlehamn, Alessandro Sette, Alan Sher, Katrin D. Mayer-Barber, Daniel L. Barber, Bongani Mayosi, Mpiko Ntsekhe, Robert J. Wilkinson, Catherine Riou

## Abstract

Our understanding of the immune response at the site of disease in extra-pulmonary tuberculosis (EPTB) is still limited. In this study, using flow cytometry, we defined the pericardial fluid (PCF) cellular composition and compared the phenotypic and functional profile of *Mycobacterium tuberculosis* (Mtb)-specific T cells between PCF and whole blood in 16 patients with pericardial TB (PCTB). We found that lymphocytes were the predominant cell type in PCF in PCTB, with a preferential influx of CD4 T cells. The frequencies of TNF-α producing myeloid cells and Mtb-specific T cells were significantly higher in PCF compared to blood. Mtb-specific CD4 T cells in PCF exhibited a distinct phenotype compared to those in blood, with greater GrB expression and lower CD27 and KLRG1 expression. We observed no difference in the production IFNγ, TNF or IL-2 by Mtb-specific CD4 T cells between the two compartments, but MIP-1β production was lower in the PCF T cells. Bacterial loads in the PCF did not relate to the phenotype or function of Mtb-specific CD4 T cells. Upon anti-tubercular treatment completion, HLA-DR, Ki-67 and GrB expression was significantly decreased, and relative IL-2 production was increased in peripheral Mtb-specific CD4 T cells. Overall, using a novel and rapid experimental approach to measure T cell response *ex vivo* at site of disease, these results provide novel insight into molecular mechanisms and immunopathology at site of TB infection of the pericardium.

## INTRODUCTION

Tuberculosis (TB) causes more deaths than any other bacterial disease. The WHO estimated that there were 1.2 million TB deaths in HIV uninfected individuals and 208 000 HIV/TB co-infected deaths in 2019 ^1^. Extra-pulmonary TB (EPTB) contributes 15% of the global TB incidence and presents diagnostic and therapeutic challenges ^1^. Meta-analyses of post-mortem studies have shown that in HIV/TB co-infected cases dissemination is frequent, with a pooled summary estimate of 87.9% of all TB cases ^3^. Importantly, 45.8% (95% CI 32.6–59.1%) of TB cases in HIV-1 infected persons were undiagnosed at time of death, highlighting the urgent need for improved rapid diagnostic tools for EPTB ^3^.

An important extra-pulmonary manifestation is pericardial TB (PCTB): the most common cause of pericardial disease in Africa, associated with debilitating complications and high mortality ^2^. PCTB disproportionately affects HIV-1 coinfected persons who are at high risk of hematogenous dissemination of TB. Although PCTB is a severe form of EPTB, our understanding of the broad range of clinicopathological PCTB phenotypes has not advanced since its first descriptions in the 1940s and 1960s ^4–6^. Moreover, studies of the immune response at the site of TB disease are limited.

Current understanding is of a spectrum of disease. At one extreme, pericardial fluid (PCF) contains a high bacillary load and represents a failure of immune control. At the other end of the spectrum, PCF can be paucibacillary with low yield of culture and polymerase chain reaction (PCR)-based tests. In this latter situation, a presumptive diagnosis of PCTB maybe made using a limited number of biomarkers with suboptimal specificity.

Despite the broad spectrum of disease phenotypes, the treatment approach is uniform and likely sub-therapeutic, as key sterilizing anti-tuberculosis drugs do not penetrate the pericardial space at sufficient concentrations to inhibit Mtb ^7^. This is of particular concern as both high bacillary load and a CD4 count <200 cells/mm^3^ in HIV-1 infected patients are predictors of mortality in PCTB ^8^. Interventions targeting both control of Mtb and HIV-1 are thus critical in HIV-1 coinfected persons with PCTB. However, regardless of ATT and anti-retroviral treatment (ART), severe complications such as re-accumulation of pericardial effusion after pericardiocentesis, compromised cardiac function due to tamponade and chronic pericardial inflammation leading to constriction (pericardial thickening with fibrosis) and death remain frequent ^9^. Thus, with few validated diagnostic, prognostic, and treatment monitoring biomarkers to guide clinicians in the management of PCTB, a better understanding of the immune response to PCTB represents an urgent unmet research priority to identify potential new clinical assessment tools ^10^.

Mtb control relies on a highly orchestrated immune response at the site of infection, and both innate and adaptive responses act synergistically to restrict Mtb growth. CD4 T cells, and in particular intact Th1 cellular responses are essential for control of Mtb ^11^. We have previously shown that a simple whole blood-based assay can be utilized to measure functional and phenotypic cellular markers that correlate with Mtb bacterial load, clinical disease severity and treatment response in PTB patients, regardless of HIV-1 status ^12, 13^.

Considering the importance of functional Mtb-specific CD4 T cells in Mtb control, in this study, we compared the frequency, polyfunctional capacity and phenotypic profile of Mtb-specific CD4 T cells in blood and PCF of patients with PCTB. We assessed whether the cellular profile associated with Mtb bacillary load and defined the effect of ATT on the peripheral Mtb-specific CD4 T cell response.

## RESULTS

### Study population

A total of 16 participants were included in the study, of which 9 were male, with a median age of 34 (Interquartile range (IQR): 28-43 years). Clinical characteristics of the study participants are listed in **Table 1** and data for each individual patient presented in **Supplemental Table 1**. The majority (87.5% [14/16]) were HIV-1 infected, and 71.4% [10/14] had either not commenced or had defaulted ART at the time of enrolment into the study. The median peripheral CD4 count of the HIV-1 infected participants were 141 cells/mm^3^ (IQR: 45-188) and the median HIV-1 viral load was 47,907 mRNA copies/ml (IQR: 1,756-178,520). All participants had large pericardial effusions with the median volume of PCF aspirated being 950 ml (IQR: 480-1200). The PCF of 9 participants (56.2%) returned Mtb culture positive results, with a median time to culture positivity of 22 days (IQR: 13-26). All aspirated effusions were exudates, with high protein content and elevated adenosine deaminase (ADA) and lactate dehydrogenase (LDH). The majority showed a predominance of mononuclear cells (median: 81.7%) over polymorphonuclear cells (median: 18.3%) by PCF cell count. The Tygerberg Diagnostic Index Score (TBH DI Score) is a weighted score incorporating clinical symptoms and laboratory values in the diagnosis of TB pericarditis (sensitivity 86%, specificity 84%)^14^. The median TBH DI Score in our study was 10, with all participants scoring above the diagnostic cut-off of 6 for TB pericarditis. No difference in any of the measured clinical parameters were observed between PCTB patients with a positive PCF Mtb culture and PCTB patients with a negative PCF Mtb culture (**Supplemental Table 2**).

**Table 1:**
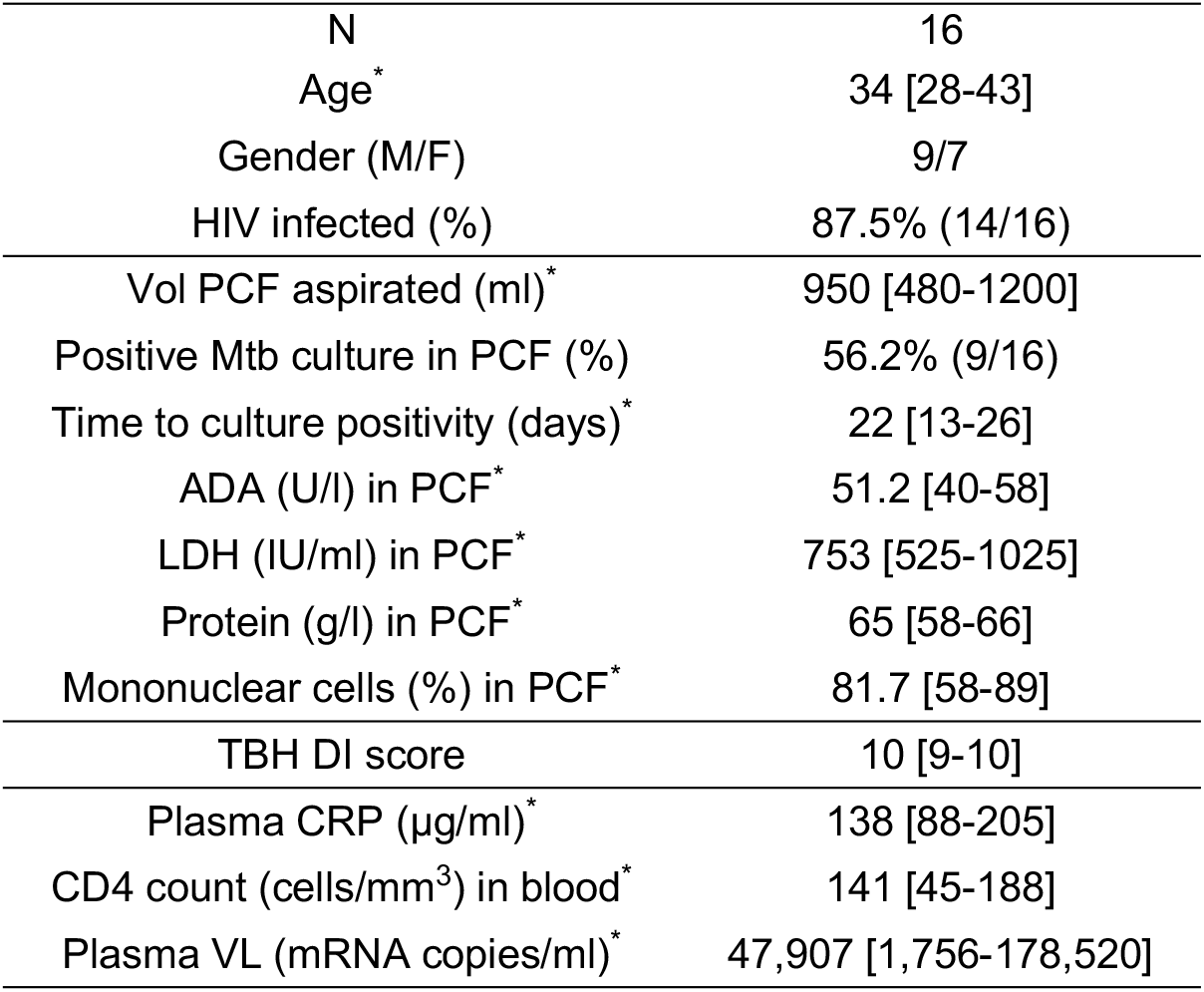
Clinical characteristic of study patients. *: Median and Interquartile range. M: Male, F: Female, PCF: Pericardial Fluid, ADA: adenosine deaminase, LDH: lactate dehydrogenase, TBH DI: Tygerberg Diagnostic Index Score (TBH DI Score) is a weighted score incorporating clinical symptoms and laboratory values in the diagnosis of TB pericarditis (sensitivity 86%, specificity 84%, ref: 14). CRP: C-reactive protein, VL: viral load.

|  |  |
| --- | --- |
| N | 16 |
| Age* | 34 [28-43] |
| Gender (M/F) | 9/7 |
| HIV infected (%) | 87.5% (14/16) |
| Vol PCF aspirated (ml)* | 950 [480-1200] |
| Positive Mtb culture in PCF (%) | 56.2% (9/16) |
| Time to culture positivity (days)* | 22 [13-26] |
| ADA (U/l) in PCF* | 51.2 [40-58] |
| LDH (IU/ml) in PCF* | 753 [525-1025] |
| Protein (g/l) in PCF* | 65 [58-66] |
| Mononuclear cells (%) in PCF* | 81.7 [58-89] |
| TBH DI score | 10 [9-10] |
| Plasma CRP (µg/ml)* | 138 [88-205] |
| CD4 count (cells/mm<sup>3</sup>) in blood* | 141 [45-188] |
| Plasma VL (mRNA copies/ml)* | 47,907 [1,756-178,520] |

### Cellular composition of pericardial fluid in PCTB

First, we examined the cellular composition of pericardial fluid (PCF) samples using flow cytometry (**Figure 1A**). As characteristically observed in PCTB ^14, 15^, we found lymphocytes to be the most abundant cell type in PCF (median 67%), while granulocytes/myeloid cells accounted for a median of 24% of the total live cell population (**Figure 1B**). However, it is notable that in six of the 16 studied participants, lymphocytes represented less than 50% of the total cell population. While not statistically significant, these participants tended to have a lower blood CD4 count and higher plasma HIV viral load compared to those with dominant lymphocytosis (median CD4: 92 vs 177 cells/mm^3^ and median HIV VL: 31,540 vs 131,600 mRNA copies/ml, respectively, data not shown). Further analysis comparing lymphocytic populations showed that the proportion of CD4 T cells was significantly higher in PCF compared to blood (median: 28.6 vs 20.8%, *P* = 0.029, respectively), while fewer CD3 negative cells were found in PCF compared to blood (median: 16.2 vs 28.5%, *P* = 0.049, respectively) (**Figure 1C**). This suggests a preferential recruitment of CD4 T cells into the pericardium during PCTB, a finding which is further supported by a significantly higher CD4/CD8 ratio in PCF compared to blood (median: 0.8 vs 0.43, *P* = 0.006, respectively) (**Figure 1D**).

**Figure 1:**
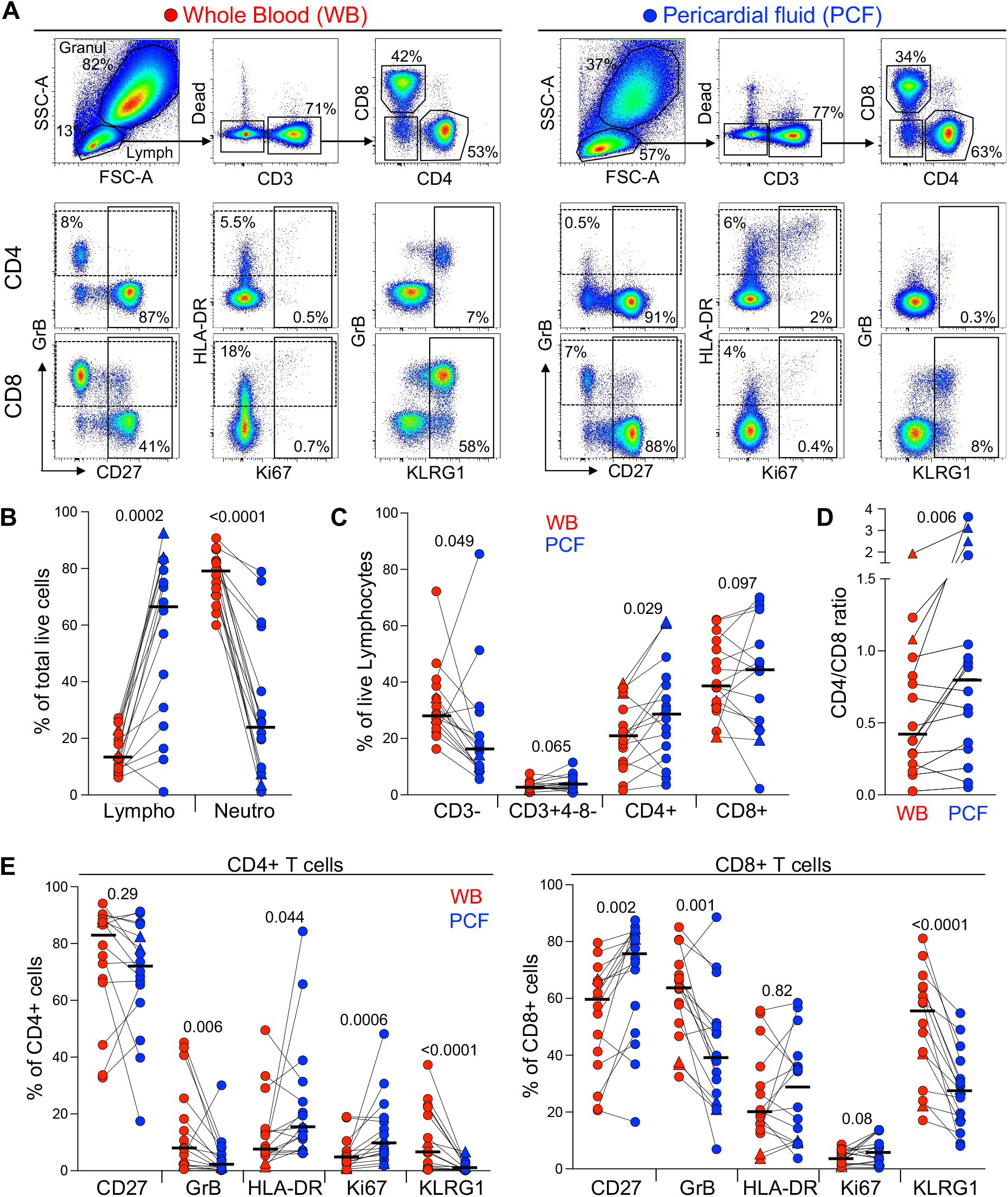
Comparison of the distribution of immune cells in whole blood and pericardial fluid (PCF) in patients with pericarditis. **(A)** Representative example of the distribution and phenotype of T cells in whole blood (red) and PCF (blue) using flow cytometry. **(B)** Proportion of lymphocyte (Lympho) and neutrophils (Neutro) defined according to their FSC/SSC profile and expressed as a percentage of total live cells in patients with pericarditis (n = 16). Triangles depict HIV-uninfected participants (n = 2). **(C)** Proportion of CD3-, CD3+CD4-CD8-, CD3+CD4+ and CD3+CD8+ cells, expressed as a percentage of total live lymphocytes. **(D)** CD4 to CD8 ratio in whole blood and PCF. **(E)** Expression of CD27, GrB, HLA-DR, Ki67 and KLRG1 in CD4 T cells (left panel) and CD8 T cells (right panel) from whole blood and PCF. Bars represent medians. Statistical comparisons were performed using a Wilcoxon rank test.

We next compared the phenotype of total CD4 and CD8 T cells between blood and PCF. CD4 T cells in PCF showed higher expression of Ki-67 and HLA-DR (median: 9.5% vs 4.4%, *P* = 0.0006 and 15.3% vs 7.4%, *P* = 0.044, respectively) and lower expression of GrB and KLRG1 (median: 1.8% vs 8%, *P* = 0.006 and 0.7% vs 6.5%, *P* < 0.0001, respectively) compared to peripheral CD4 T cells (**Figure 1E**, left panel). Similarly, CD8 T cells in PCF expressed less GrB and KLRG1 compared to peripheral CD8 T cells (median: 38.9% vs 63.5%, *P* = 0.001 and 27.4% vs 54.8%, *P* < 0.0001, respectively). Moreover, in PCF, CD8 T cells showed higher expression of CD27 compared to blood (median: 75.8% vs 59.5%, *P* = 0.002) (**Figure 1E**, right panel). The lower expression of KLRG1 on T cells found at the disease site is likely related to the poor ability of these cells to migrate to site of infection, as previously demonstrated in murine models ^16, 17^. Overall, these results suggest that infiltrating CD4 T cells were more activated and exhibited a more differentiated memory profile compared to peripheral cells, while CD8 T cells in PCF had a lower cytotoxic potential and displayed features of an earlier differentiated phenotype compared to their blood counterparts.

### Comparison of the Mtb-specific Immune response in paired blood and PCF

To measure the Mtb-specific immune response, we used an approach where *ex vivo* unprocessed whole blood or pericardial fluid samples were freshly stimulated for a short time (5h) using an Mtb peptide pool combined with g-irradiated Mtb (H37Rv) in the presence of protein transport inhibitors (added at the onset of the stimulation). The combination of peptides and whole bacteria allows the simultaneous detection of the Mtb-specific T cell response and toll like receptor (TLR)-dependent innate responses. Moreover, this type of assay has the advantage of preserving all cell subsets and soluble components present *in vivo*, thus maintaining a more physiologic environment ^18^.

First, although the flow cytometry panel used in this study was not designed to measure the innate immune response in detail, we were able to compare Mtb-induced production of TNF-α by peripheral and PCF granulocytes/myeloid cells (**Figure 2A**). Figure 2B shows that while Mtb stimulation led to TNF-α production in granulocytes/myeloid cells from both compartments, the frequencies of TNF-α producing cells was ~10 fold higher [IQR: 3.3 - 20.2] at the site of TB disease compared to blood (median: 1.1% vs 10.2%, respectively, *P* < 0.0001). TNF-α production was observed almost exclusively in the HLA-DR+ population, suggesting that Mtb-responding cells are likely classical (CD14++CD16-) or intermediate (CD14++CD16+) monocytes ^19^. No difference in the frequency of TNF-α responding granulocytes/myeloid cells in PCF was observed between patients with a positive or a negative PCF Mtb culture (**Supp Figure 1A**).

**Figure 2:**
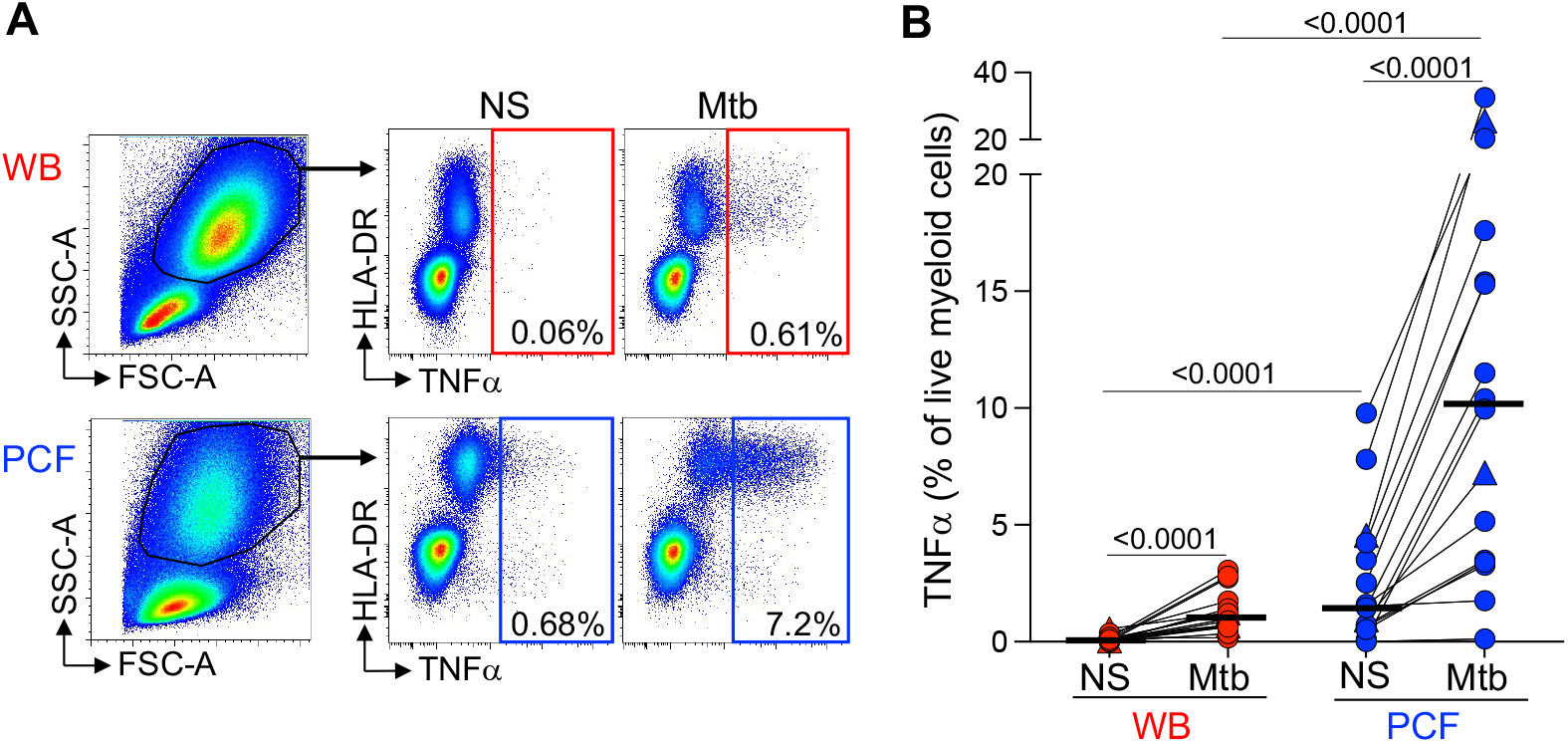
Comparison of TNFα production by granulocytes/myeloid cells in response to Mtb stimulation between whole blood and PCF. **(A)** Representative example of TNFα-producing cells in responses to Mtb antigen stimulation. Granulocytes/myeloid cells were identified based on their FCS/SSC profile. **(B)** Frequency of TNFα-producing granulocytes/myeloid cells in unstimulated (Uns) and Mtb-stimulated samples (Mtb). The triangles depict HIV-uninfected participants (n = 2). Bars represent medians. Statistical comparisons were performed using a Wilcoxon rank test for paired samples or a Mann-Whitney test for unpaired samples.

Next, to compare the adaptive immune response between blood and PCF, we measured IFN-γ, TNF-α and IL-2 expression in CD4 T cells in response to Mtb antigens (**Figure 3A**). Only one participant had no detectable Mtb-specific CD4 response in blood. The frequency of Mtb-specific CD4 T cells detected in PCF was significantly higher (~ 5-fold) than in blood for all measured cytokines (**Figure 3B**). There was no correlation between the frequency of Mtb-specific CD4 T cells in blood and PCF *(P* = 0.59, r = 0.14, **Figure 3C**). Moreover, we did not find any association between the frequency of Mtb-specific CD4 T cells and the number of absolute CD4 T cells in blood, nor between the frequencies of Mtb-specific CD4+ T cells and the percentage of total CD4 T cells in PCF (*P* = 0.11, r = −0.40 and *P* = 0.32, r = −0.26, respectively). Despite the major difference in magnitude of the Mtb-specific CD4 response between compartments, we did not observe any significant differences in polyfunctional capacity (i.e. expression of IFN-γ, TNF-α or IL-2) of Mtb-specific CD4 T cells between blood and PCF. Mtb-specific CD4 T cells exhibited a highly polyfunctional profile with a sizable proportion (median: 58% for blood and 55% for PCF) of cells co-expressing IFN-γ, TNF-α and IL-2 (**Figure 3D**). Additionally, Mtb culture positivity in PCF did not associate with the magnitude or polyfunctional capacities of Mtb-specific CD4 T cells in blood or in PCF (**Supp Figure 1B&C**).

**Figure 3:**
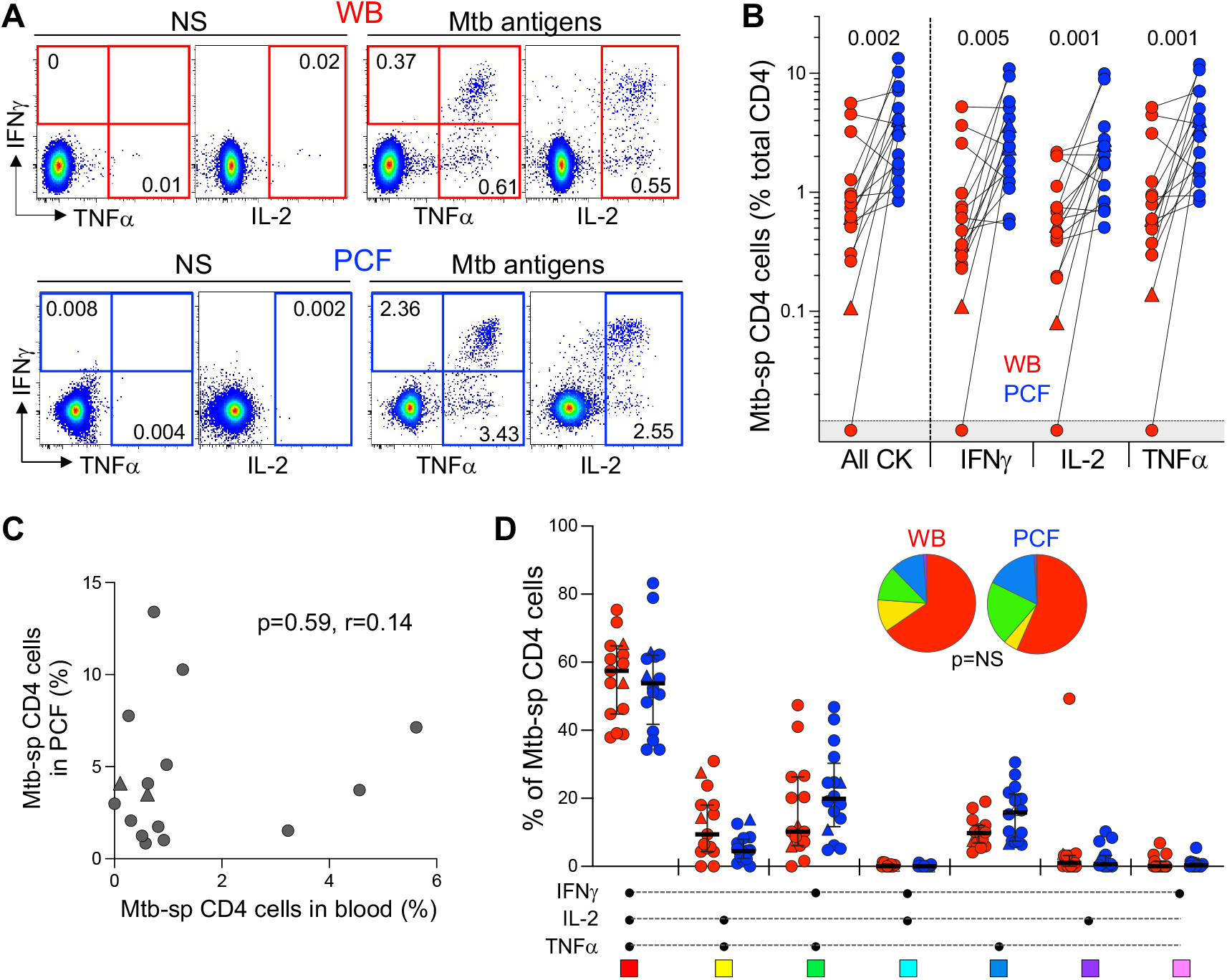
Comparison of the functional profile of Mtb-specific CD4 T cells between whole blood and PCF. **(A)** Representative example of IFNγ, IL-2, and TNFα production by CD4 T cells in unstimulated (unstim) and Mtb-stimulated whole blood and PCF samples. Number represents cytokine positive cells expressed as a percentage of total CD4 T cells. **(B)** Magnitude of Mtb-specific CD4 T cells in whole blood (red) and PCF (blue). The triangles depict HIV-uninfected participants (n = 2). Bars represent medians. Statistical comparisons were defined using a Wilcoxon rank test. **(C)** Correlation between the frequency of Mtb-specific CD4 T cells in blood and at site of disease (PCF). The triangles depict HIV-uninfected participants (n = 2). Correlation was tested by a two-tailed non-parametric Spearman rank test. **(D)** Polyfunctional profile of Mtb-specific CD4 T cells in whole blood (red) and PCF (blue). The x-axis displays all possible cytokine combinations (flavors), the composition of which is denoted with a dot for the presence of IL-2, IFNγ and TNFα. The proportion of each flavor contributing to the total Mtb-specific CD4 response per individual is shown. The median (black bar) and interquartile ranges (box) are shown. Each flavor is color-coded, and data are summarized in the pie charts, representing the median contribution of each flavor to the total Mtb response. No statistically significant differences were observed using a Wilcoxon rank test to compare response patterns between groups and a permutation test to compare pies.

Detailed phenotyping was performed to define and compare the activation and maturation profile of Mtb-specific CD4 T cells from blood and PCF. Firstly, irrespective of the studied compartments, Mtb-specific CD4 T cells exhibited high expression of HLA-DR (ranging from 48 to 97%) and Ki67 (ranging from 3.6 to 79%) and low expression of CD27 (ranging from 9 to 46%); profiles characteristic of active TB disease, as previously reported ^20–23^ (**Figure 4A**). Paired comparison of blood and PCF revealed that Mtb-specific CD4 T cells in PCF had distinct characteristics compared to their peripheral counterparts, with elevated expression of GrB (medians: 36.6% vs 16%, respectively, *P* = 0.017) and lower expression of MIP-1β (36.6% vs 16%, *P* = 0.003), CD27 (16.2% vs 20%, *P* = 0.03) and KLRG1 (0.5% vs 3.8%, *P* = 0.002) (**Figure 4A**). However, despite differences in the level of expression of GrB and KLRG-1 between blood and PCF, the phenotype of Mtb-specific CD4 T cell in the blood associated with the profile of PCF Mtb-specific CD4 T cells for GrB, KLRG-1, CD153, Ki-67, and HLA-DR, with the strongest correlations being observed for GrB (*P* =0.0034, r = 0.74) and CD153 (*P* = 0.0034, r = 0.74) (**Figure 4B**). Additionally, the phenotypic profile of PCF Mtb-specific CD4 T cells was comparable regardless of PCF culture status (**Figure 4C**).

**Figure 4:**
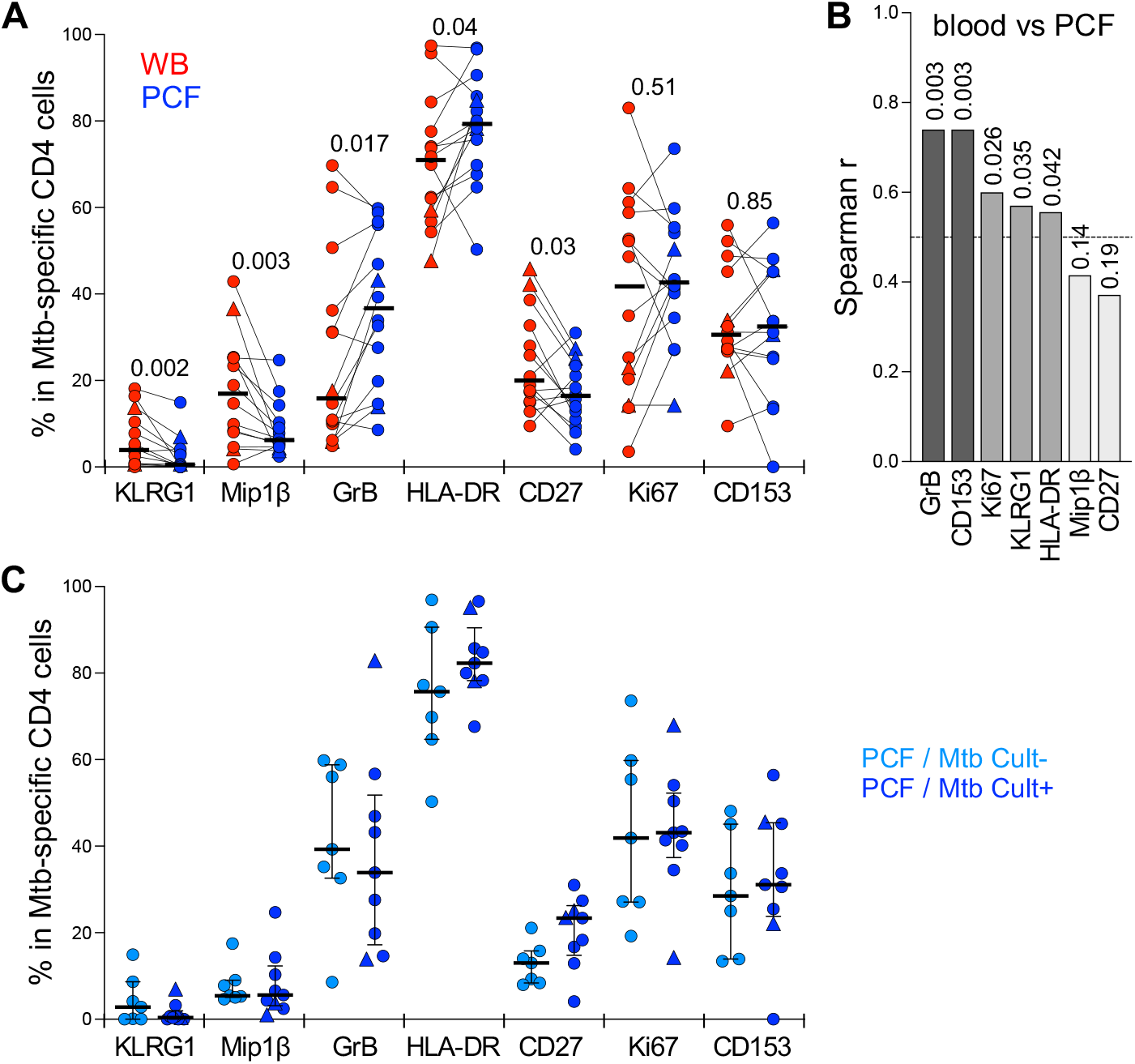
Comparison of the phenotypic profile of Mtb-specific CD4 T cells in blood and PCF. **(A)** Expression of GrB, CD27, HLA-DR, Ki67, KLRG1, CD153 and MIP1β in Mtb-specific CD4 T cells (i.e. producing any measured cytokine IFNγ, IL-2, and TNFα). Only paired samples with Mtb-specific responses >30 events are depicted (n=14). Bars represent medians. The triangles depict HIV-uninfected participants (n = 2). Statistical comparisons were performed using a Wilcoxon rank test. **(B)** Spearman correlation r values between indicated Mtb-specific CD4 T cell phenotype in blood and PCF. *P* values are indicated for each comparison. **(C)** Comparison of the phenotypic profile of Mtb-specific CD4 cells from PCF between patients who are PCF Mtb culture negative (PCF/Mtb Cult-, light blue, n=7) or PCF Mtb culture positive (PCF/Mtb Cult+, dark blue, n=9). Bars represent medians. Statistical comparisons were performed using the Mann-Whitney test.

We also investigated the frequency and phenotype of Mtb-specific CD8 T cells in these participants (**Supp Figure 2**). As previously described in TB patients ^24^, a CD8 T cell response was not observed in all participants. Mtb-specific CD8 T cells were detected in blood from 5 out of 16 patients (32%), while 8 of 16 patients (50%) had a detectable CD8 T cell response in PCF (**Supp Figure 2B**). As in the case of CD4 responses, we did not find any phenotypic differences between blood and PCF CD8 responses (**Supp Figure 2C**). Of note, compared to Mtb-specific CD4 T cells, CD8 T cells exhibited limited capacity to produce IL-2, expressed significantly higher level of GrB and Mip-1β; with CD153 expression being undetectable (**Supp Figure 2D**). The activation profile (defined with HLA-DR and Ki67 expression) were comparable between Mtb-specific CD4 and CD8 T cell responses in PCF and blood (**Supp Figure 2D**).

### Impact of ATT on blood Mtb-specific CD4 T cell responses

In the context of pericardial TB (and extra-pulmonary TB, in general), the monitoring of the response to treatment is particularly challenging due to the lack of sensitive and specific tools. In this study, we defined the impact of ATT on the frequency, polyfunctional and phenotypic profile of peripheral Mtb-specific CD4 T cells pre- and post-ATT (at week 24 or 52, depending on sample availability). While successful ATT did not alter the magnitude of Mtb-responding CD4 T cells (**Figure 5A**), it significantly modified their functional capacity, with the proportion of IL-2 and TNF-α dual producing CD4 T cells becoming significantly expanded post-ATT, counterbalanced by a decrease of IFN-γ and TNF-α dual producing CD4 T cells (**Figure 5B**). More importantly, ATT led to major changes in the phenotype of the peripheral Mtb-specific CD4 response (**Figure 5C&D**). Post-ATT, HLA-DR, Ki67 and GrB expression by Mtb-specific CD4 T cells were significantly reduced compared to pre-treatment (medians: 15.7 vs 70% for HLA-DR; 2.8 vs 35% for Ki67 and 0.9 vs 10.9% for GrB, respectively), while no changes were observed for CD153, CD27 or Mip-1β expression (**Figure 5D**). Overall, these results suggest that assessing the activation profile of Mtb-specific CD4 T cells could aid monitoring of treatment response in extra-pulmonary TB.

**Figure 5:**
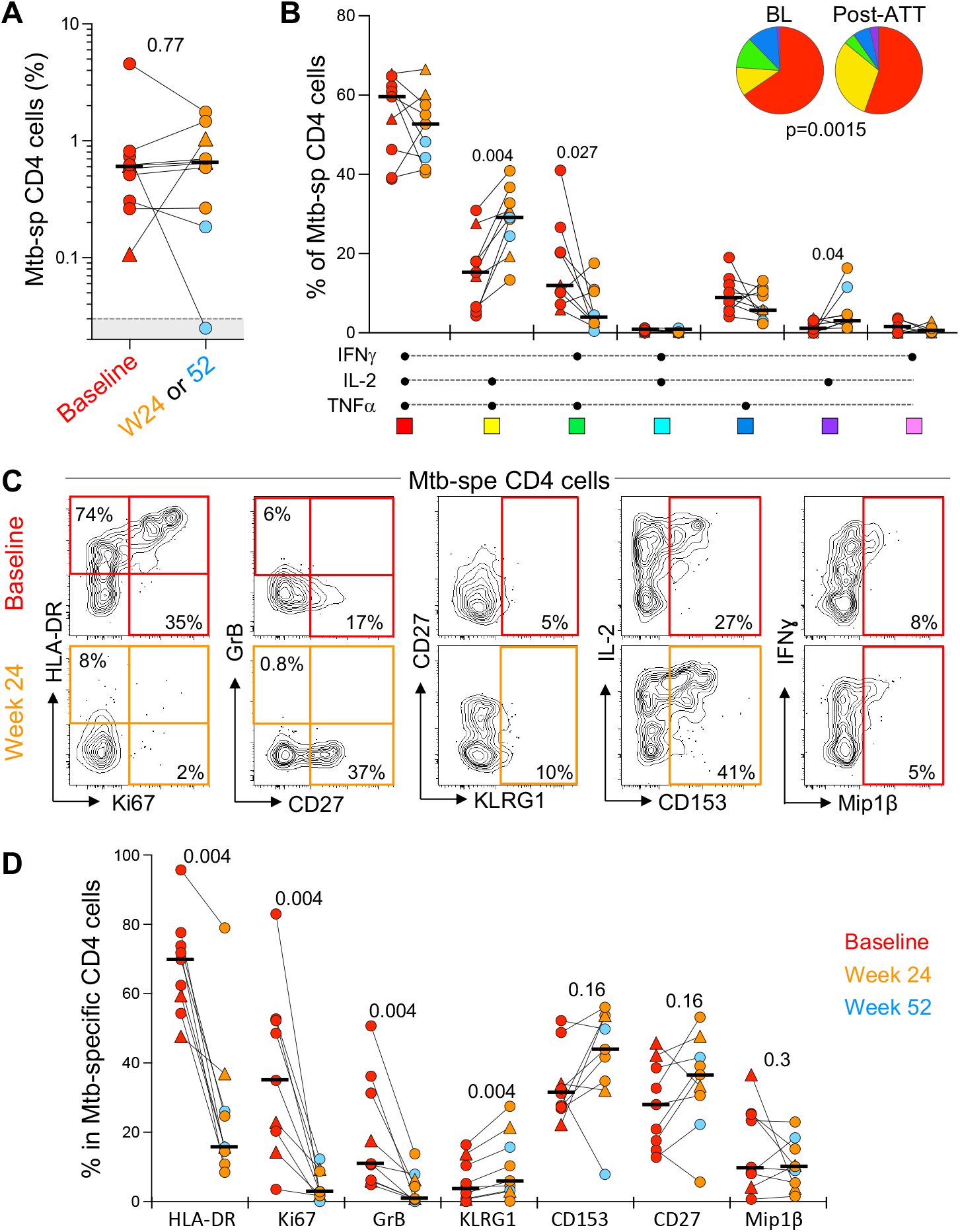
Evolution of the magnitude, polyfunctional capacity and phenotypic profile of Mtb-specific CD4 T cells response in whole blood from baseline (pre-ATT initiation) to ATT completion (24 or 52 weeks post-ATT). **(A)** Frequency of Mtb-specific CD4 T cells pre-ATT (Baseline, red) and post ATT (24 weeks, orange or 54 weeks, light blue). The triangles depict HIV-uninfected participants (n = 2). Bars represent medians. Statistical comparisons were performed using the Mann-Whitney test. **(B)** Polyfunctional profile of Mtb-specific CD4 T cells pre-ATT (BL) and post ATT (W24 or W52). The median and interquartile ranges are shown. A Wilcoxon rank test was used to compare cytokine combination between groups and a permutation test to compare pies. **(C)** Representative flow plots of HLA-DR, Ki67, GrB, CD27, KLRG1, CD153 and MIP1β expression in Mtb-specific CD4 T cells pre- and post-ATT in one patient. **(D)** Summary graph of the expression of phenotypic markers measured. Only paired samples are depicted (n=9). Red symbols correspond to Baseline samples, orange to W24 samples and blue to W52 samples. The triangles depict HIV-uninfected participants (n = 2). Bars represent medians. Statistical comparisons were performed using the Wilcoxon rank test.

## DISCUSSION

Pericardial TB is an understudied and severe form of extra-pulmonary TB ^10^. Only two previous published studies have used flow cytometry to assess the cellular profile of PCF in PCTB ^15, 25^. Here, we demonstrate that a simple laboratory methodology (derived from a previously described whole blood assay ^13^) can successfully be applied to PCF, revealing differences and similarities between the two compartments. Using this approach, we investigated the overall cellular profile of PCF, assessed whether PCF Mtb culture positivity associate with the Mtb specific cellular response and evaluated the evolution of peripheral Mtb-specific CD4 T cell responses in relation to ATT.

In accordance with previous reports, we found a significantly higher frequency of lymphocytes in PCF compared to blood, whereas neutrophils predominated in blood ^14^. Additionally, we observed significantly higher frequencies of CD4 T cells in PCF compared to blood, whereas preferential recruitment was not observed for CD8 T cells. This effect is likely to be even more pronounced in cohorts comprised of higher numbers of HIV-1 uninfected participants, given the findings by Reuter *et al*. showing lower frequencies of CD4 T cells and higher frequencies of CD8 T cells in PCF of HIV-1 co-infected compared to HIV-1 uninfected patients with PCTB ^15^. Here, we report that Mtb-specific CD4 T cells are more abundant in PCF than in blood in HIV-1 infected individuals, similar to what has been reported in studies of other TB disease sites (e.g. lymph nodes, broncho-alveolar lavage and pleural fluid) ^26–31^. A previous study comparing PCF and blood of PCTB patients only observed increased PCF CD4 T cell responses in HIV-1 uninfected individuals through examining numbers of spot forming cells in response to ESAT-6 detected by IFN-γ ELISpot ^25^. This difference is likely attributable to the use of different laboratory assays (ELISpot versus flow cytometry), sample type (cryopreserved cells versus *ex vivo* whole blood or whole pericardial fluid) and stimuli (ESAT-6/CFP-10 versus Mtb300). Despite markedly higher frequencies of cytokine producing Mtb-specific CD4 T cells in PCF compared to blood, we did not observe any significant differences in their functional capacity, in keeping with previous reports ^26, 32^. Similarly, in a prior study, we did not observe significant differences in cytokine production between ESAT-6/CFP-10 specific CD4 T cells in PCF and blood of pericardial patients ^25^. The frequency of Th1 cytokine producing Mtb-specific CD4 T cells may play an important role for Mtb control, as demonstrated by a study involving low dose Mtb infection of non-human primates which compared T cell cytokine production at granuloma level in the lung to peripheral blood ^33^. They observed significant variability in cytokine production between different granulomas in the same animal, with sterile granulomas comprising a higher frequency of cytokine (IL-17, TNF and any Th1 cytokine [IFN-γ, IL-2, or TNF]) producing T cells than non-sterile granulomas. It is thus likely that the predominance of Mtb-specific CD4 T cells in PCF over blood reflects homing of these cells to the pericardium to attempt contribution to Mtb control.

We hypothesized that the presence of a higher Mtb load, as denoted by positive Mtb culture status, would enhance T cell proliferation (thus frequency), and may promote further differentiation and activation of responsive T cells ^20, 34^. However, we did not observe any major differences in the Mtb-specific T cell profile according to PCF Mtb culture status. It should be borne in mind that PCF Mtb culture is of poor sensitivity to diagnose PCTB ^10^. Nonetheless, this finding warrants further investigation, especially since time to culture positivity in PCF significantly predicts mortality in PCTB and considering the important established role of an efficient Th1 response in protecting against Mtb ^8^. Future studies of pericardial biopsy or pericardiectomy samples could potentially shed further light on this by analyzing T cell responses and Mtb load at the granuloma level.

Growing evidence points towards the necessity to elicit a balanced immune response to Mtb, with the predominance of either pro- or anti-inflammatory mediators being detrimental in the containment and elimination of Mtb ^33, 35^. IFN-γ and TNF-α are examples of essential pro-inflammatory cytokines that are required in the immune response to Mtb ^36^, especially through their capacity to activate infected macrophages to eliminate Mtb ^37^, but in excess, these cytokines may also exacerbate immunopathology ^38–40^. Excessive TNF-α can induce mitochondrial reactive oxygen species that lead to necrotic cell death and activate matrix metalloproteinases both of which have been associated with lung tissue damage during PTB ^39, 41^. We found PCF granulocytes to have significantly higher capacity to produce TNF-α in response to Mtb compared to blood granulocytes. Whilst we found no significant difference in frequency of Mtb-specific granulocytes producing TNF-α in culture positive compared to negative PCF, it remains to be investigated whether higher frequencies of TNF-α producing granulocytes are associated with enhanced immune pathology and complications such as pericardial fibrosis.

The clinical management of pericardial TB is complicated owing to the lack of sensitive and specific diagnostic and treatment monitoring tools. A definitive diagnosis of PCTB relies on the detection of Mtb in PCF or in pericardial tissue, and thus requires invasive sampling. The sensitivity of conventional PCF culture techniques is between 50 - 65%, whilst Xpert MTB/RIF of PCF has 66 - 66.7% sensitivity when assessed against a composite reference ^42, 43^. Whilst Xpert MTB/RIF yields rapid results, Mtb culture can take up to 4-6 weeks to yield results. The diagnosis thus rests on the presence of a lymphocyte predominant effusion with elevated levels of the biomarker ADA. Measurement of IFN-γ in PCF and the use of IFN-γ release assays (IGRAs) have produced variable, but promising results as diagnostic tests, but is currently limited to the research setting pending further optimization, head-to-head comparison with current composite reference standards and studies evaluating its scalability in low resource settings ^44, 45^. Few, if any, advances have recently been made in efforts to monitor treatment efficacy in EPTB. Indeed, PCTB treatment response is only assessed clinically owing to the impracticality of invasive sampling of the pericardium, contrasting with PTB where follow up sputums can be obtained to monitor Mtb clearance. We and others have previously shown that a simple, whole blood-based assay assessing the activation profile of Mtb-specific CD4 T cells can be used as a non-sputum-based method to track both disease severity and response to ATT ^13, 20, 21, 46, 47^. Here, we build on these findings showing that 1) in PCTB patients, the activation profile of peripheral Mtb-specific CD4 T cells is comparable to those observed in active PTB and 2) successfully treated PCTB is associated with significantly decreased expression of HLA-DR, Ki-67 and GrB and expansion of IL-2 producing Mtb-specific CD4 T cells compared to baseline. HLA-DR has been shown to be a robust biomarker to discriminate latent TB infection from PTB ^13, 20, 21, 47^ and EPTB ^23^, and to monitor PTB treatment response ^13, 20, 46, 47^. Here, we show that HLA-DR expression on peripheral Mtb-specific CD4 T cells significantly decreases between baseline and completion of ATT in PCTB, thus extending the potential utility of this biomarker for the monitoring of treatment response in PCTB.

Our study was limited by small numbers of HIV-1 uninfected participants. However, we have previously shown that our panel of phenotypic and functional T cell markers perform equally well regardless of HIV-1 status ^13^. Moreover, this study would have benefited from the inclusion of patients with non-tuberculous pericardial effusion to further ascertain the specificity of HLA-DR expression on Mtb-specific T cells as potential TB diagnostic biomarkers. In this study, our objective was not to investigate immune correlates of adverse outcomes such as pericardial constriction, other complications or mortality, with these remaining important research priorities for future studies. The performance of multiple PCF cultures, or the additional testing of Xpert MTB/RIF with reporting of cycle threshold values may have increased the sensitivity of our estimate of Mtb burden.

Nonetheless, in view of the dearth of knowledge on immune responses at TB disease site, our study demonstrates a novel and rapid experimental approach to measure T cell response *ex vivo* at site of disease. This technique allowed us to define key differences and similarities between the Mtb immune response in PCF and blood and may aid to the diagnostic and/or treatment monitoring of PCTB patients. The latter will be of particular importance as pharmacodynamic readout in ongoing trials aiming to optimize treatment regimens for PCTB, with the current standard of care ATT lacking sufficient concentrations in the pericardial space to inhibit Mtb ^7^.

## Author’s contributions

C.R., E.d.B., D.L.B., K.D. M-B, M.N., B.M. and R.J.W. designed the study. P.H., A.J. and E.d.B. recruited the study participants. S.R., E.d.B. and C.R. performed the whole blood and PCF assay. E.d.B. and C.R. performed the flow experiments. C.R. analysed and interpreted the data. A.Se. and C.S.L.A. provided critical reagents. R.J.W., A.Sh. and C.R. obtained funding to support the project. C.R. and E.d.B. wrote the manuscript with all authors contributing to providing critical feedback.

## METHODS

### Study population

Participants with suspected PCTB recruited from the Groote Schuur Hospital Cardiology Unit. Only adults (≥ 18 years of age) who were undergoing pericardiocentesis as part of the routine management of their pericardial effusion and who had received no more than 3 doses of ATT prior were included. Pregnancy, severe anemia (Hemoglobin ≤ 7g/dL), multi-drug resistant TB and severe concurrent opportunistic infection were exclusion criteria. All participants provided written informed consent and the study was approved by the University of Cape Town Human Research Ethics Committee (HREC: 050/2015, DMID protocol 15-0047). At the time of pericardiocentesis the study team collected PCF and paired blood samples for analysis. Only participants with definite (Mtb culture positive) or probable PCTB were included in the study. Probable PCTB was defined per criteria from Mayosi *et al*. where there was evidence of pericarditis with microbiologic confirmation of Mtb-infection elsewhere in the body and/or an exudative, lymphocyte predominant pericardial effusion with elevated adenosine deaminase (≥35 U/L) ^48^. Study participants were followed up over the course of their ATT and up to one year.

### Pericardial fluid, blood collection and stimulation assay

Pericardial fluid was obtained at the time of pericardiocentesis, placed in sterile Falcon tubes and transported to the laboratory at 4°C. Blood was collected in sodium heparin tubes at the time of pericardiocentesis. Both blood and pericardial fluid were processed within 3 hours of collection. The whole blood or whole PCF assay were adapted from the protocol described by Hanekom *et al*. ^49^. Briefly, 0.5 ml of whole blood or 1 ml of whole PCF were stimulated with a pool of 300 Mtb-derived peptides (Mtb300, 2 μg/ml) ^50^ combined with g-irradiated Mtb (H37Rv, 100 μg/ml, obtained through BEI Resources, NIAID, NIH) at 37°C for 5 hours in the presence of the co-stimulatory antibodies, anti-CD28 and anti-CD49d (1 μg/ml each; BD Biosciences, San Jose, CA, USA) and Brefeldin-A (5 μg/ml; Sigma-Aldrich, St Louis, MO, USA) and Monensin (5 μg/ml, BD Biosciences, San Jose, CA, USA). The combination of Mtb peptides and inactivated whole bacteria allows the simultaneous detection of Mtb-specific T cell response and toll like receptor (TLR)-dependent innate response. Unstimulated cells were incubated with co-stimulatory antibodies and Brefeldin-A only. Red blood cells were then lysed in a 150 mM NH_4_Cl, 10 mM KHCO_3_, 1 mM Na_4_EDTA solution. Cells were then stained with a Live/Dead Near-InfraRed dye (Invitrogen, Carlsbad, CA, USA) and then fixed using a Transcription Factor Fixation buffer (eBioscience, San Diego, CA, USA), cryopreserved in freezing media (50% fetal bovine serum, 40% RPMI and 10% dimethyl sulfoxide) and stored in liquid nitrogen until use.

### Cell staining and flow cytometry

Cryopreserved cells were thawed, washed and permeabilized with a Transcription Factor perm/wash buffer (eBioscience). Cells were then stained at room temperature for 45 minutes with the following antibodies: CD3 BV650 (OKT3; Biolegend, San Diego, CA, USA), CD4 BV785 (OKT4; Biolegend), CD8 BV510 (RPA-T8; Biolegend), CD27 PE-Cy5 (1A4CD27; Beckman Coulter, Brea, CA, USA), HLA-DR BV605 (L243; Biolegend), Ki67 PerCPcy5.5. (B56, BD), Granzyme B (GrB) BV421 (GB11, BD), Killer cell Lectin-like Receptor G1 (KLRG1) APC (13F12F2, eBioscience), IFNγ BV711 (4S.B3; Biolegend), TNFα PEcy7 (Mab11; Biolegend), IL-2 PE/Dazzle (MQ1-17H12, Biolegend), Mip-1β Alexa Fluor 488 (#24006, R&D systems, Minneapolis, MN, USA) and CD153 (R&D116614, R&D). Samples were acquired on a BD LSR-II and analyzed using FlowJo (v9.9.6, FlowJo LCC, Ashland, OR, USA). The granulocytes/monocytes population was defined based on their FSC/SSC characteristics. A positive cytokine response was defined as at least twice the background of unstimulated cells. To define the phenotype of Mtb300-specific cells, a cut-off of 30 events was used.

### Statistical analyses

Statistical tests were performed in Prism (v9.1.2; GraphPad, San Diego, CA, USA). Nonparametric tests were used for all comparisons. The Kruskal-Wallis test with Dunn’s Multiple Comparison test was used for multiple comparisons and the Mann-Whitney and Wilcoxon matched pairs test for unmatched and paired samples, respectively.

## SUPPLEMENTARY MATERIAL

**Supp Table 1:**
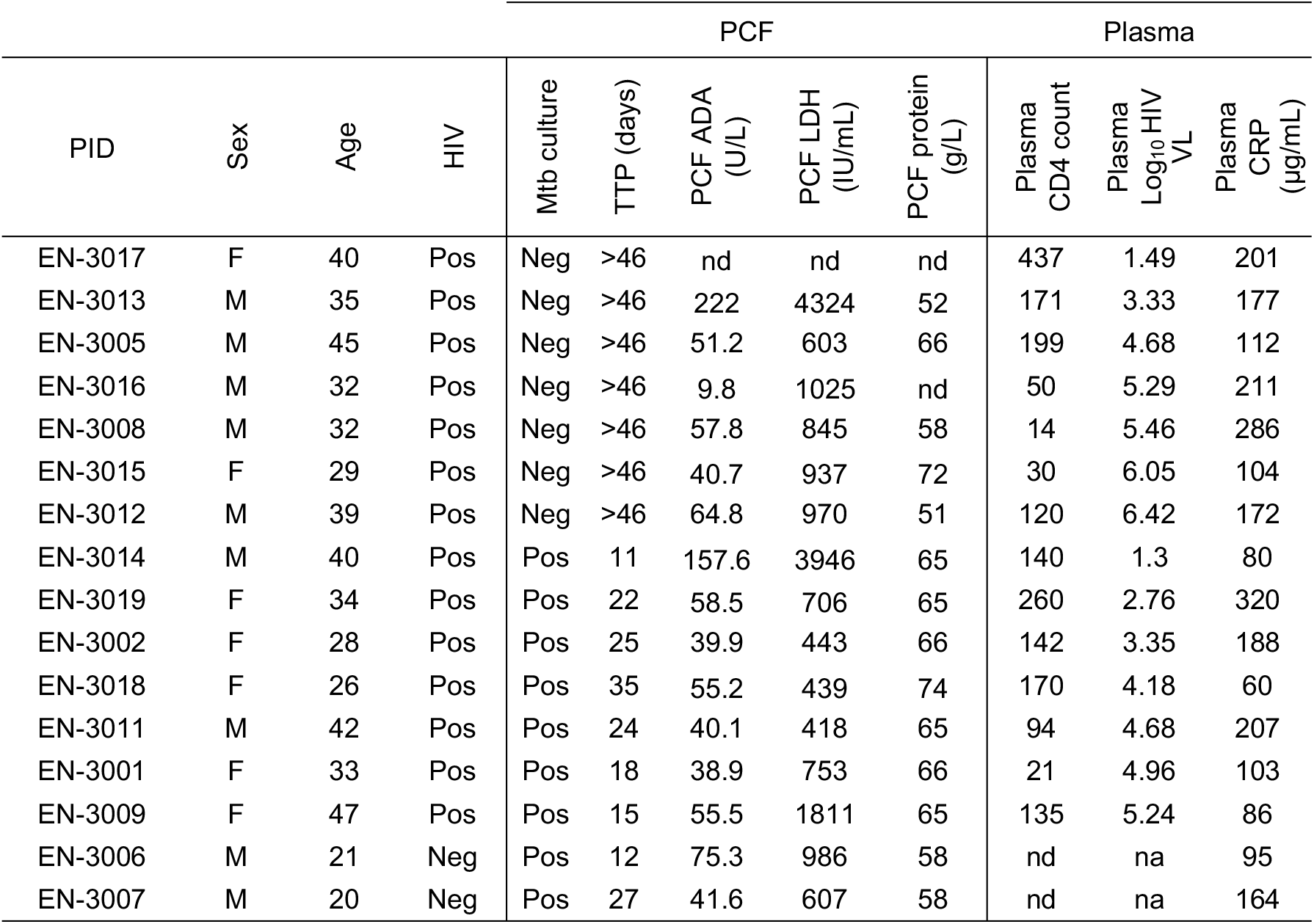
Clinical characteristics in each studied patient (n=16). PCF: Pericardial Fluid, M: Male, F: Female, TTP: time to positivity, ADA: adenosine deaminase, LDH: lactate dehydrogenase. CRP: C-reactive protein, VL: viral load. Nd: not done, na: not applicable.

| PID | Sex | Age | HIV | PCF |  |  |  |  | Plasma |  |  |
| --- | --- | --- | --- | --- | --- | --- | --- | --- | --- | --- | --- |
|  |  |  |  | Mtb culture | TTP (days) | PCF ADA (U/L) | PCF LDH (IU/mL) | PCF protein (g/L) | Plasma CD4 count | Plasma Log<sub>10</sub> HIV VL | Plasma CRP (µg/mL) |
| EN-3017 | F | 40 | Pos | Neg | >46 | nd | nd | nd | 437 | 1.49 | 201 |
| EN-3013 | M | 35 | Pos | Neg | >46 | 222 | 4324 | 52 | 171 | 3.33 | 177 |
| EN-3005 | M | 45 | Pos | Neg | >46 | 51.2 | 603 | 66 | 199 | 4.68 | 112 |
| EN-3016 | M | 32 | Pos | Neg | >46 | 9.8 | 1025 | nd | 50 | 5.29 | 211 |
| EN-3008 | M | 32 | Pos | Neg | >46 | 57.8 | 845 | 58 | 14 | 5.46 | 286 |
| EN-3015 | F | 29 | Pos | Neg | >46 | 40.7 | 937 | 72 | 30 | 6.05 | 104 |
| EN-3012 | M | 39 | Pos | Neg | >46 | 64.8 | 970 | 51 | 120 | 6.42 | 172 |
| EN-3014 | M | 40 | Pos | Pos | 11 | 157.6 | 3946 | 65 | 140 | 1.3 | 80 |
| EN-3019 | F | 34 | Pos | Pos | 22 | 58.5 | 706 | 65 | 260 | 2.76 | 320 |
| EN-3002 | F | 28 | Pos | Pos | 25 | 39.9 | 443 | 66 | 142 | 3.35 | 188 |
| EN-3018 | F | 26 | Pos | Pos | 35 | 55.2 | 439 | 74 | 170 | 4.18 | 60 |
| EN-3011 | M | 42 | Pos | Pos | 24 | 40.1 | 418 | 65 | 94 | 4.68 | 207 |
| EN-3001 | F | 33 | Pos | Pos | 18 | 38.9 | 753 | 66 | 21 | 4.96 | 103 |
| EN-3009 | F | 47 | Pos | Pos | 15 | 55.5 | 1811 | 65 | 135 | 5.24 | 86 |
| EN-3006 | M | 21 | Neg | Pos | 12 | 75.3 | 986 | 58 | nd | na | 95 |
| EN-3007 | M | 20 | Neg | Pos | 27 | 41.6 | 607 | 58 | nd | na | 164 |

**Supp Table 2:** Comparison of the clinical characteristics between patients with a PCF positive Mtb culture (n=9) and patients with a PCF negative Mtb culture (n=7). *: Medians and Interquartile ranges. M: Male, F: Female, PCF: Pericardial Fluid, ADA: adenosine deaminase, LDH: lactate dehydrogenase, TBH DI score: Tygerberg Diagnostic Index Score is a weighted score incorporating clinical symptoms and laboratory values in the diagnosis of TB pericarditis (sensitivity 86%, specificity 84%, ref: 14). CRP: C-reactive protein, VL: viral load. Statistical comparisons were performed using the Mann-Whitney test.

|  | Positive Mtb cult<br>in PCF | Negative Mtb cult<br>in PCF | *P* values |
| --- | --- | --- | --- |
| N | 9 | 7 |  |
| Age* | 33 [24-45] | 35 [32-40] | 0.84 |
| Gender (M/F) | 4/5 | 5/2 | na |
| HIV positive (%) | 77.8% (7/9) | 100% (7/7) | na |
| Vol PCF aspirated (ml)* | 1040 [455-1275] | 835 [460-1140] | 0.71 |
| Time to culture positivity (days)* | 22 [13-26] | na | na |
| ADA (U/l) in PCF* | 55 [40-68] | 46 [31-99] | 0.60 |
| LDH (IU/ml) in PCF* | 301 [202-525] | 324 [226-432] | 0.46 |
| Protein (g/l) in PCF* | 65 [61-66] | 64 [55-69] | 0.59 |
| Mononuclear cells (%) in PCF* | 81.7 [48-89] | 82.3 [71-88] | 0.74 |
| TBH DI score* | 10 [8.5-10] | 10 [8.5-10] | 0.91 |
| Plasma CRP (µg/ml)* | 103 [83-197] | 177 [104-211] | 0.41 |
| CD4 count (cells/mm<sup>3</sup>) in blood* | 140 [94-170] | 171 [30-199] | 0.80 |
| Plasma VL (mRNA copies/ml)* | 15,223 [569-90,452] | 109,051 [2,151-287,630] | 0.21 |

**Supp Figure 1:**
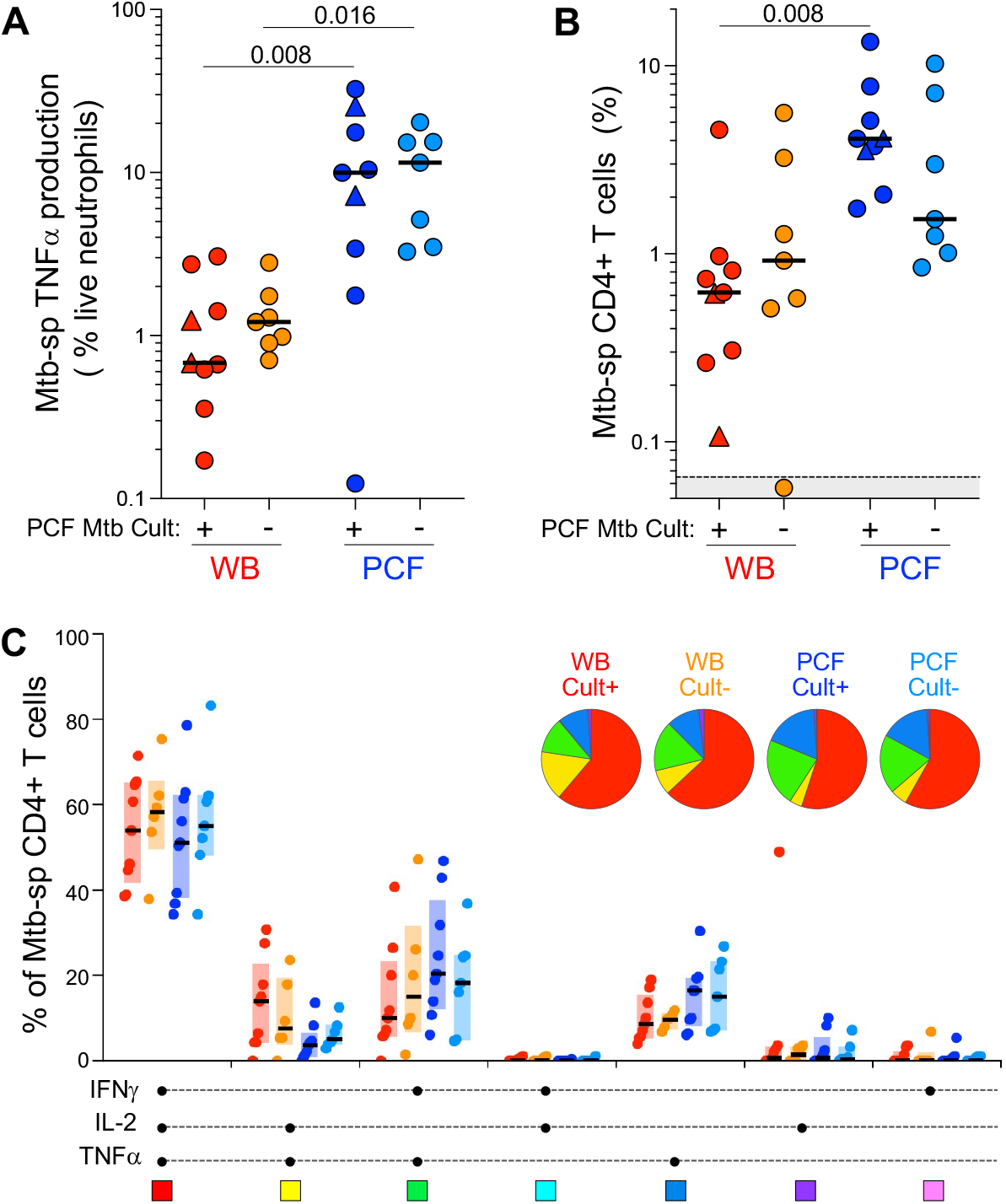
Comparison of the frequency and polyfunctional profile of immune responses in blood and PCF, stratify based on PCF Mtb culture status. (A) Frequency of TNFα+ Neutrophils in response to Mtb stimulation in whole blood and PCF from PCF Mtb culture negative (n = 7) and PCF Mtb culture positive (n = 9) patients. (B) Frequency of Mtb-specific CD4+ T cells in whole blood and PCF from patients who are PCF Mtb culture negative or PCF Mtb culture positive. (C) Polyfunctional profile of Mtb-specific CD4+ T cells. Statistical comparisons were performed using the Wilcoxon rank test for paired samples or the Mann-Whitney test for unpaired samples.

**Supp Figure 2:**
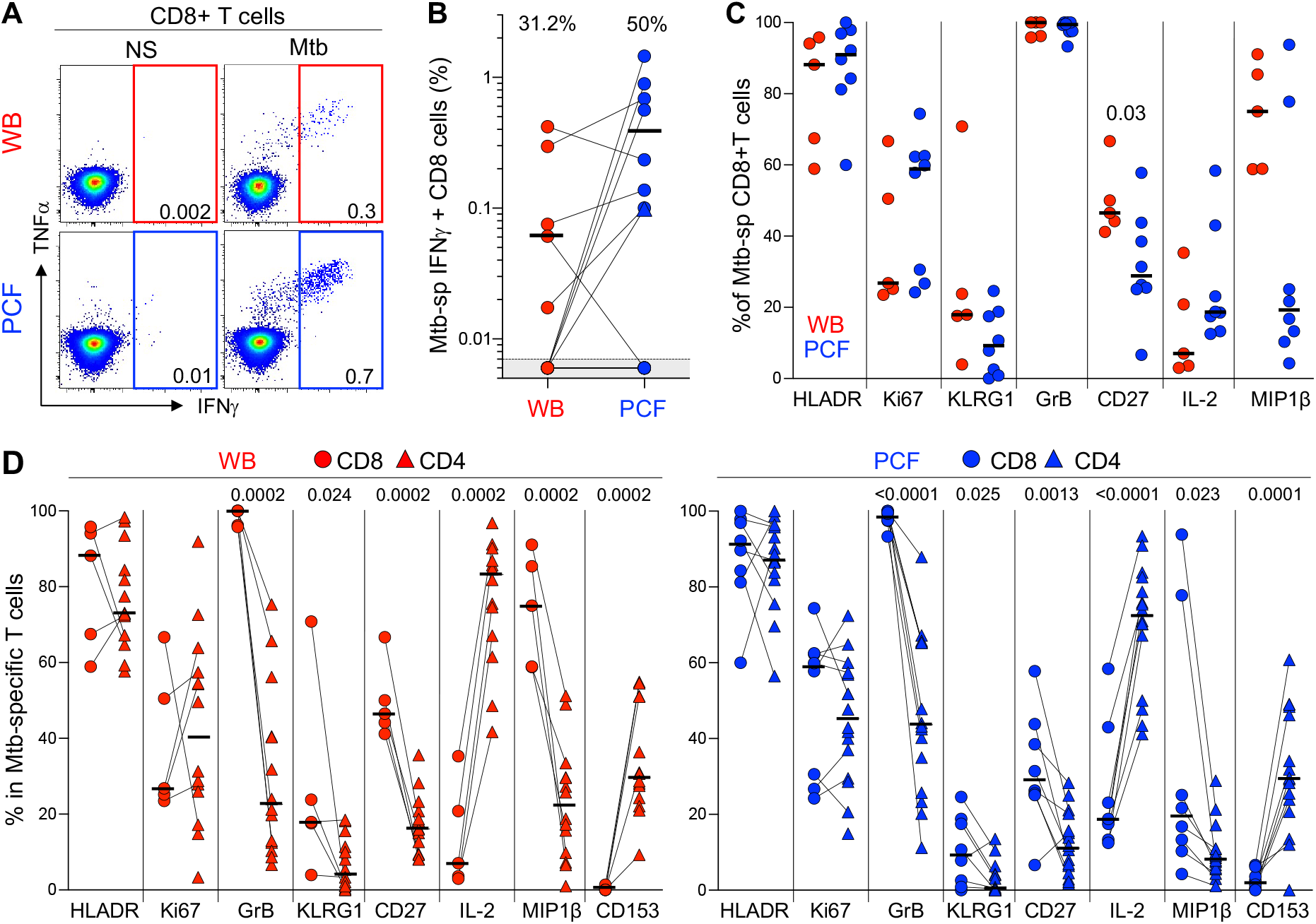
Frequency and phenotypic profile of Mtb-specific CD8+ T cell response in Blood and PCF. (A) Representative example of flow cytometry plots showing the expression of TNFα and IFNγ in CD8+ T cells from blood (top) and PCF (bottom) in one PCTB patient. (B) Comparison of the frequency of Mtb-specific CD8+ T cells in whole blood and PCF (n = 16). The proportion of participant exhibiting a detectable Mtb-specific CD8+ T cell response is indicated at the top of the graph. Bars represent medians of responders. (C) Comparison of the phenotypic profile of Mtb-specific CD8+ T cells in whole blood (n = 5) and PCF (n = 8). (D) Comparison of the phenotypic profile of Mtb-specific CD4+ and CD8+ T cells in whole blood (left panel and PCF (right panel). Bars represent medians. Statistical comparisons, performed using a Mann-Whitney test, are reported at the top of the graph.

